# Genome-wide SNPs reveal the social structure and invasion pathways of the invasive tropical fire ant (*Solenopsis geminata*)

**DOI:** 10.1101/2022.07.20.500883

**Authors:** P Lenancker, T Walsh, S Metcalfe, D Gotzek, BD Hoffmann, L Lach, WT Tay, S Elfekih

## Abstract

Elucidating invasion pathways of invasive species is often challenging because invasive populations frequently have low genetic diversity caused by genetic bottlenecks during introduction events. Genome-wide sequencing such as Restriction Site-Associated DNA Sequencing (RADseq) can overcome these challenges by generating thousands of genome-wide single nucleotide polymorphic (SNP) markers. The tropical fire ant, *Solenopsis geminata*, is a global invader with low genetic diversity in its introduced range, making RADseq one of the best available methods to investigate its population genetics. We used double digest RADseq to generate 3,834 SNPs to compare the genetic diversity of *S. geminata* in its introduced range to its most likely source of introduction, determined the invasion pathways among populations at an unprecedented level of detail for this species, and determined the social structure of *S. geminata* workers collected in 13 locations worldwide. We found that introduced *S. geminata* went through a strong genetic bottleneck. We also identified multiple secondary introduction events among *S. geminata* populations, indicating that the bridgehead effect is an important driver in the global spread of this species. We found that all colonies in the introduced range were polygyne (i.e., with more than one queen) which may increase their invasion success and potential to cause adverse effects.

## INTRODUCTION

Retracing the origin and pathways of biological invasions is crucial to identify main sources of introduction, potential management strategies (e.g., targeted surveillance, biological control agents), and study fundamental concepts of invasion ecology (Dlugosch & Parker, 2008; Hulme, 2009; Wilson et al., 2009). For example, determining whether propagule pressure originates from a single (e.g., *Batillaria attramentaria*: Asian mud snail, Miura et al., 2006) or multiple source populations (e.g., *Dreissena polymorpha*: zebra mussel, Stepien et al., 2005) may enable the development of targeted biosecurity surveillance strategies (Colunga-Garcia et al., 2013). Additionally, identifying the native range of an introduced species can be used to determine whether it has lost genetic diversity and/or undergone adaptive evolution following its introduction (Dlugosch & Parker, 2008). Unfortunately, the development of international trade makes it increasingly difficult to determine main invasion pathways. Identifying the native range of historically introduced invaders can also be challenging. For example, some organisms (e.g., fouling molluscs, algae) have been unintentionally transported among continents since the 15th century, making the introduction history of these organisms difficult to retrace without the use of genomic tools (Carlton, 1999).

Introduced populations commonly go through a genetic bottleneck during their establishment which results in a low level of genetic diversity in these populations. For example, the allelic richness of introduced *Lasioglossum leucozonium* (solitary sweat bee) populations has been reduced by 76% compared to the native populations’, and only three microsatellite multilocus genotypes have been found among introduced populations of *Aegilops triuncialis* (Barbed goatgrass) vs. 57 in their native range (Meimberg et al., 2006; Zayed et al., 2007). As a result, DNA markers commonly used for genetic assessments such as microsatellite markers and mitochondrial DNA gene sequences may have low variability in their invasive range and provide limited information on main invasion pathways. For example, low genetic variation at microsatellite loci in introduced populations of *Codium fragile* (green alga) only enabled large scale invasion patterns to be drawn, and the low variability at mitochondrial DNA gene sequences in native and invasive populations of *Oryctes rhinoceros* (coconut rhinoceros beetle) prevented resolution of its invasion pathways (Provan et al., 2005; Reil et al., 2016).

The development of next-generation technologies such as Restriction site-Associated DNA Sequencing (RADseq) in the past decade has facilitated the study of invasive species. RADseq is a tool capable of generating hundreds to millions of genome-wide SNP markers, providing a reduced representation of the genome without needing a reference genome (Andrews et al., 2016). The refining of laboratory protocols and RADseq library preparations makes it possible to use small amounts of DNA (< 100ng) for sequencing (Andrews et al., 2016; Nygaard & Wurm, 2015). RADseq is a powerful tool used to address critical scientific questions such as population structure, gene flow events, species boundaries, and colonisation routes of invasive pests (Eaton & Ree, 2013; Elfekih et al., 2018, 2021, 2022; Nadeau et al., 2014; Sherpa & Després, 2021; Storer et al., 2017). RADseq has also considerably improved our ability to identify the introduction routes and origin of invasive populations with low genetic diversity. For example, the introduction history of *Sargassum muticum* (brown seaweed) was successfully reconstructed with RADseq loci whereas microsatellite markers failed to detect genetic variation across its introduced range (Le Cam et al., 2020). Another RADseq study inferred global routes of introduction of *Rattus norvegicus* (brown rat), whereas a previous study using mitochondrial markers identified their centre of origin but was unable to resolve relationships among invasive populations (Puckett et al., 2016; Song et al., 2014).

Despite being particularly useful to retrace the invasion history of invaders with low genetic diversity such as ants, RADseq has never been used to retrace global pathways of introduced ants to the best of our knowledge. Over 200 ant species are known to have established outside their native range with 31 species considered to have major environmental, economic, health, and social impacts (Angulo et al., 2022; Bertelsmeier et al., 2017; Gruber et al., 2022; Holway et al., 2002). Studies on the introduction history of global ant invaders using microsatellite markers and/or mitochondrial DNA gene sequences have all identified several main introduction pathways, but some population movements, especially between introduced populations, could not be reconstructed due to the low variability of the markers used in the ants’ introduced range (Ascunce et al., 2011; Foucaud et al., 2010; Vogel et al., 2010). For example, the mean microsatellite allelic richness of *Linepithema humile* (Argentine ant, Vogel et al., 2010) in its introduced range was 33% lower than in its native range and a large majority (311/322) of unique mtDNA haplotypes were exclusively found in native populations of *Solenopsis invicta* (red imported fire ant, Ascunce et al., 2011). Secondary introductions are a major driver accelerating global ant invasions through the bridgehead effect in which introduced populations act as the source of additional invasions (Bertelsmeier et al., 2018). Identifying fine scale movements between introduced populations is therefore crucial to develop a better understanding of ant invasion pathways, and RADseq may solve some of the shortcomings of markers traditionally used in these studies. Data generated with RADseq methods can also be used to answer questions about colony structure and population dynamics (e.g., unicoloniality, social structure) which may affect ants’ invasiveness and impact potential (Holway et al., 2002; Tsutsui & Suarez, 2003). For example, polygyne (multiple queened) populations of invasive ants can reach higher densities than monogyne (single-queened) ones, which contributes to their ability to dominate ecosystems (Holway et al., 2002; Macom & Porter, 1996; Tsutsui & Suarez, 2003).

*Solenopsis geminata* (tropical fire ant) is one of the world’s six most widespread, abundant, and damaging invasive ants and is established globally throughout tropical regions (Gruber et al., 2022; Hodgson & Clarke, 2014; Holway et al., 2002; Wetterer, 2011). Colonies of *S. geminata* can be either monogyne or polygyne (Holway et al., 2002). Both monogyne and polygyne colonies appear to be present in its native and introduced range (Adams et al., 1976; Lenancker et al., 2019; Mackay et al., 1990; Nipitwattanaphon et al., 2020; Ross et al., 2003; Trible et al., 2018; Wauters et al., 2018; Williams & Whelan, 1991) but, the invasive population established in Taiwan appears to be exclusively monogynous (Lai et al., 2015). A genomics study of *S. geminata* global populations using microsatellite and mitochondrial markers concluded that the ant was likely transported from southwestern Mexico to the Philippines in the 16^th^ century and, from there to the rest of the Old World (Gotzek et al., 2015). However, finer scale movements among introduced populations could not be determined due to the low variability of microsatellite and mitochondrial markers across the species’ introduced range. Four mtDNA haplotypes and an average (±SD) of 4±2.7 alleles per microsatellite markers were identified in introduced populations (Gotzek et al., 2015). Because RADseq studies have successfully reconstructed population movements which could not be detected with microsatellite and mitochondrial markers, it is a suitable method to investigate secondary invasion patterns for a species with low genetic diversity in its introduced populations such as *S. geminata* as well as the social structure of colonies (Gotzek et al., 2015; Le Cam et al., 2020; Lenancker et al., 2019; Puckett et al., 2016; Song et al., 2014).

We used ddRADseq (double digest RADseq, Peterson et al. 2012), a variant method of RADseq, to generate genome-wide SNPs and provide the most detailed investigation yet into the invasion pathways, potential bottleneck, and social structure of *S. geminata* at a global scale. We used specimens collected in southwestern Mexico in the native range of *S. geminata* as it is the most likely source for all introduced populations and 12 locations in its introduced range, including three locations (Cambodia, Malaysia, and Ashmore Reef), for which its introduction history had not previously been investigated (Gotzek et al., 2015). Our objectives were to (i) compare the genetic diversity of *S. geminata* in its introduced range to its most likely source of introduction, (ii) determine the invasion pathways among populations at an unprecedented level of detail for this species, and (iii) determine the social structure of *S. geminata* in its introduced range.

## METHODS

### Solenopsis geminata samples

We genetically analysed 177 *S. geminata* workers belonging to 28 colonies from 13 distinct geographic locations including three locations (Cambodia, Malaysia, and Ashmore Reef) which had not previously been investigated. We used a single worker per colony or 3 to 8 workers per colony (mean ± SD: 7±1) depending on the availability of the material (Figure 1, Table 1). Most samples were from introduced populations in the Indo-Pacific region. For the native range, we only included specimens from southwestern Mexico, which is the most-likely source of the introduced populations (Gotzek et al., 2015). Workers were preserved in 80% ethanol and stored at 4°C until molecular analysis.

**Table 1.**
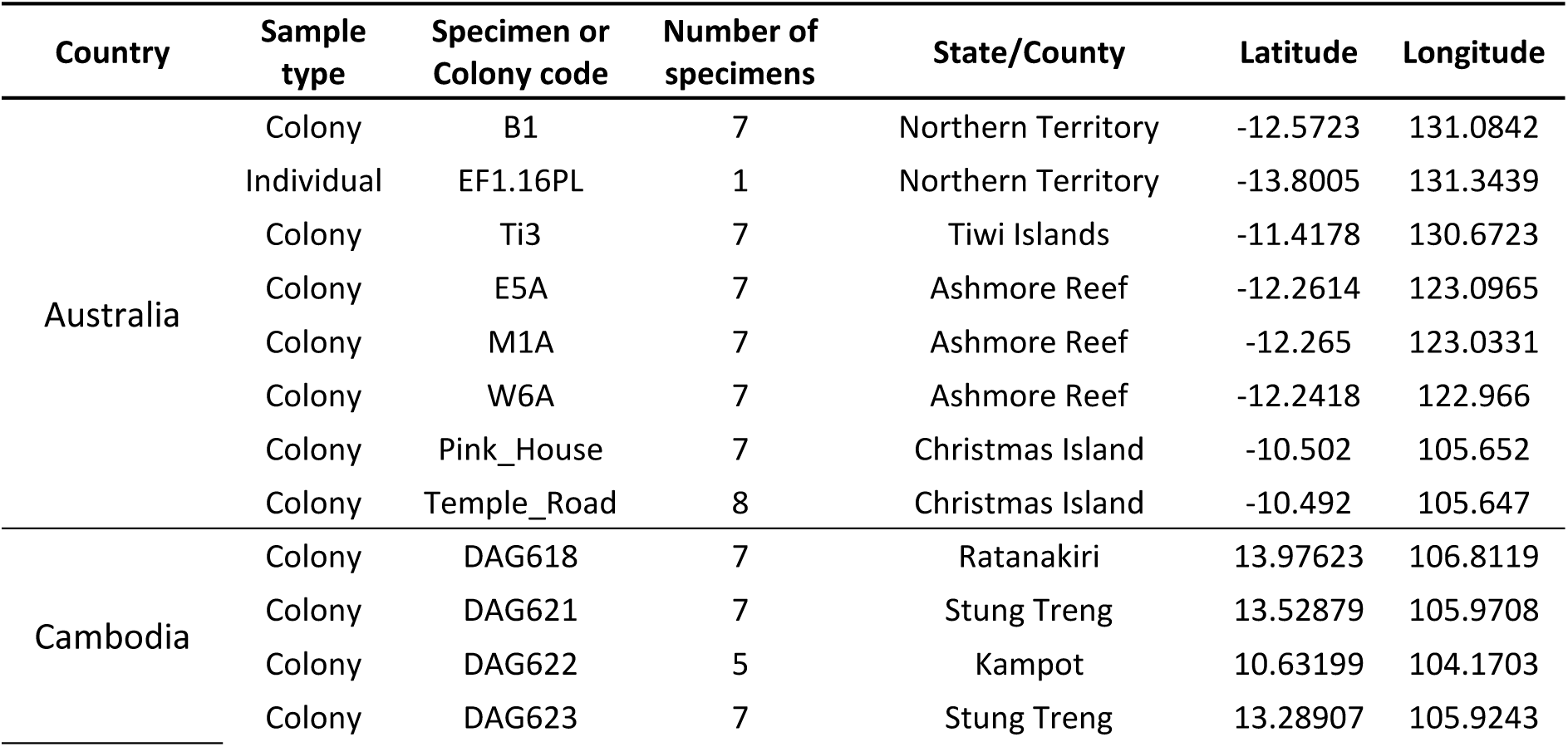
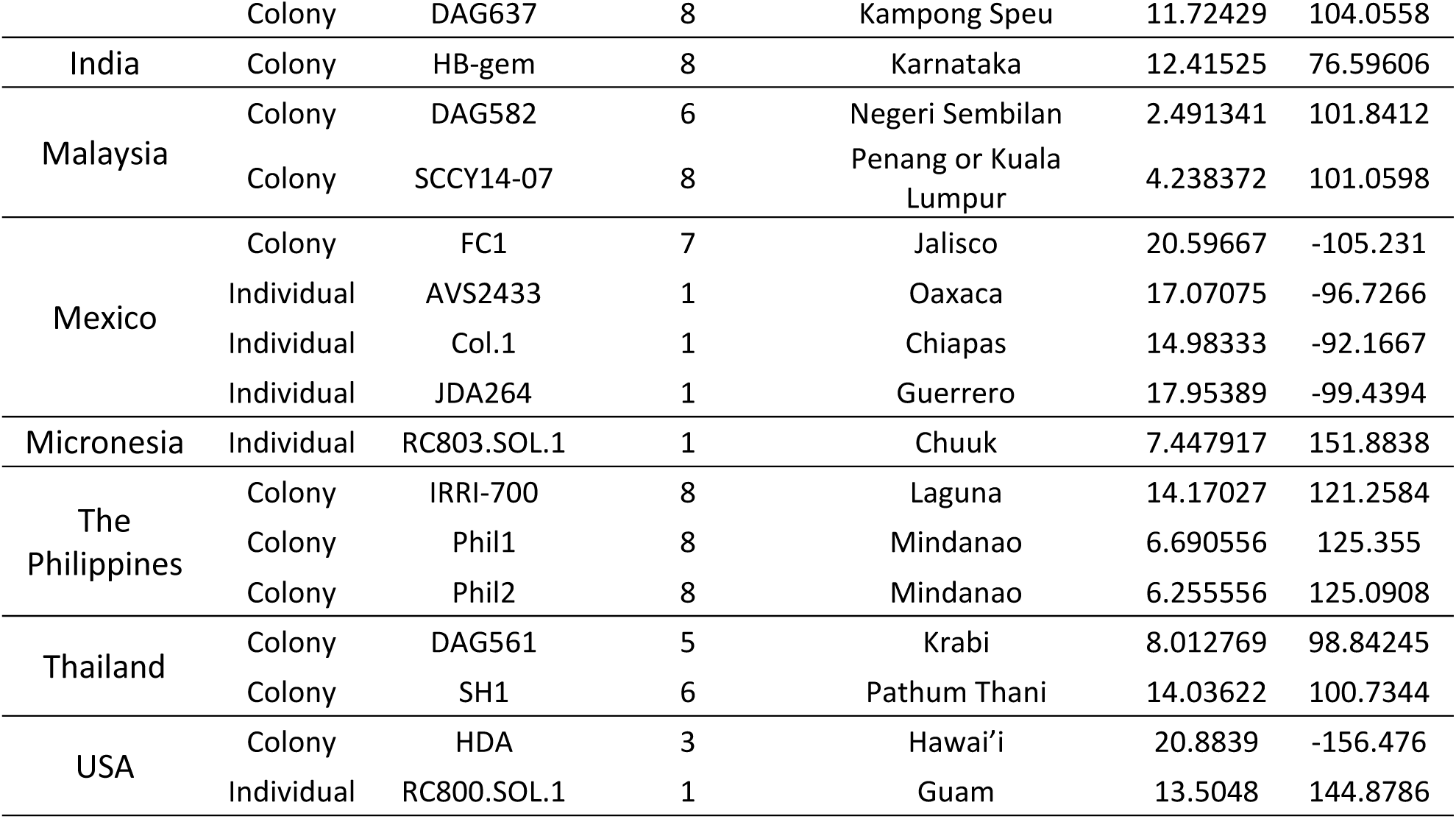
Final sample set of 157 *Solenopsis geminata* specimens which passed the data filtering steps (see SNP calling and filtering section in the methods) and were used in this study

**Figure 1.**
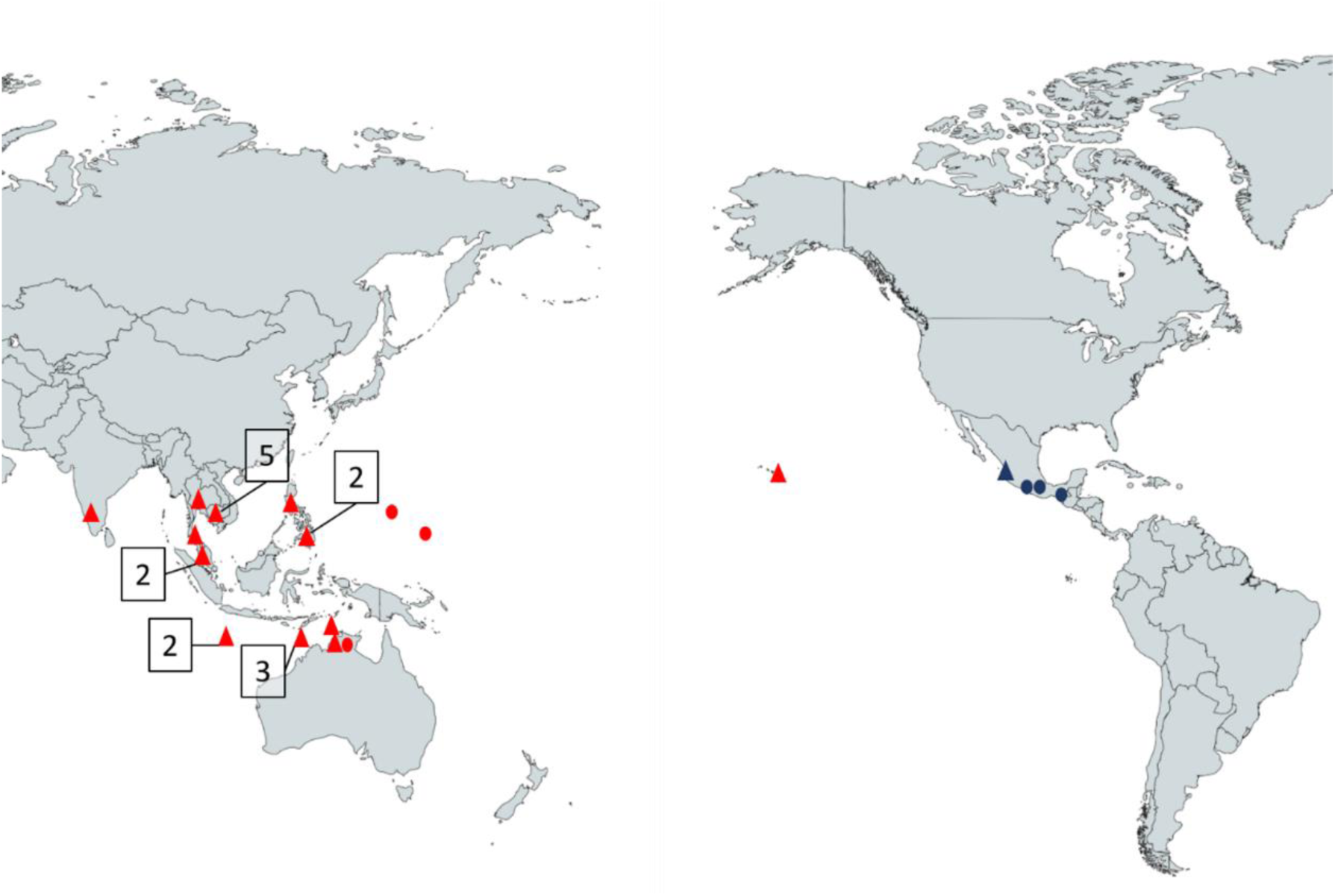
Collection locations of *S. geminata* specimens analysed in this study. Red: specimens collected in the introduced range. Blue: specimens collected in the native range. Circle: single specimen. Triangle: multiple specimens from a colony. Numbers show the number of colonies sampled if more than one originated from this location.

### Genomic DNA extraction

Individual workers were placed in a 1.5 ml Eppendorf® tube, snapped frozen in liquid nitrogen and homogenised with a sterile pestle. We extracted total genomic DNA (gDNA) using the Qiagen DNeasy® Blood and Tissue kit modifying the manufacturer’s instructions slightly (half the lysis buffer volumes and eluted the gDNA in 20µl Elution Buffer containing 10mM Tris-HCL). We quantified gDNA with a Qubit® 3.0 fluorometer (Life Technologies, Eugene, Oregon, USA) and selected gDNA samples with the highest concentrations.

### ddRAD library preparation

The ddRADseq protocol was adapted from Peterson et al. (2012) and DaCosta and Sorenson (2014). Briefly, we prepared double-digest of 50-100ng of gDNA with 20 units each of high-fidelity restriction enzymes, *PstI* and *EcoRI* (New England Biolabs, Ipswich MA, USA), in CutSmart buffer in a final volume of 50µL. Samples were digested in a thermocycler for 30 min at 37°C, then heat inactivated for 20 min at 65°C and slowly cooled to room temperature and held at 4°C. We assessed and quantified the fragment size distribution on the 2100 BioAnalyzer (Agilent, CA, USA) using the High Sensitivity DNA chips. We ligated 50µL of each digested gDNA sample to uniquely barcoded P1 and P2 adapters in a 70µL reaction containing 4µL of 50 nM P1 adapter, 12µL of 50 nM P2 adapter, 1µL of 2000 units of T4 DNA ligase (New England Biolabs, Ipswich MA, USA), 0.6µL of 0.15 mM rATP (Promega, Madison, WI, USA), 2µL of 10x NEBuffer 2 and 0.4µL of water. We ligated the samples on a thermal cycler for 30 min at 22°C, then heat inactivated for 20 mins at 65°C, and slowly cooled to room temperature and held at 4°C. Five µL of ligated gDNA from each sample were combined to create a pool of individuals. The samples were size selected with 0.65x volume of Agencourt® AMPure® XP magnetic bead solution (Beckman Coulter, Brea, CA, USA), and suspended in 20µL elution buffer (Qiagen, Hilden, Germany). We prepared the final library amplification in a 60µL PCR reaction with 15µL of size selected DNA, 30µL of Phusion High Fidelity 2x Master Mix (New England Biolabs, Ipswich MA, USA), 3µL of 10 µM each of P1 and P2 primers (Table S1) and 9µL water. We ran the samples at 98°C for 30 s, followed by 20 cycles of 98°C for 10 s, 60°C for 30 s and 72°C for 40 s, with the final elongation at 72°C for 5 min. We cleaned the PCR product with 1.8x volume of Agencourt® AMPure® XP magnetic bead solution (Beckman Coulter, Brea, CA, USA), and suspended in 40µL elution buffer (Qiagen, Hilden, Germany). We quantified the ddRAD libraries on the Qubit 3.0 fluorometer (Life Technologies, Eugene, Oregon, USA) and assessed the size fragments on a Tape Station 2200 (Agilent, CA, USA) using the High-Sensitivity DNA screen tapes. We normalized the final library to 4nM for sequencing on the HiSeq platform (Illumina, CA, USA).

### SNP calling and filtering

We used FastQC version 0.11.8 (Andrew, 2010) to assess the quality of the raw fastq sequences. We dropped ten low-quality samples based on the FastQC reports, and trimmed the remaining sequences by quality (Phred quality score higher than 20) to a length of 50bp using Trimmomatic version 0.38 (Bolger et al., 2014).

We conducted SNP calling using a *de novo* approach on the 164 samples that passed the data filtering step. We used the software pipeline stacks version 2.2 (JM Catchen & Amores, 2011; Julian Catchen et al., 2013) for SNP calling. We first used ustacks to align our short-read sequences into stacks with a minimum depth of coverage of three (3X). We used the final coverage information given at the end of the ustacks step to eliminate samples (n=7) with less than 50% reads. We then used ctsacks on the 157 remaining samples to build a catalogue of consensus loci with 2 mismatches allowed between sample loci. We used sstacks to search the putative loci built by ustacks against the catalog produced by ctsacks.

We then used tsv2bam to orientate the data by locus and gstacks to align reads to the locus and call SNPs. We used the last stacks pipeline step, populations, to generate population genetic statistics and test three loci filtering scenarios. The minimum percentage of individual required to process a locus I for our population was conducted for three threshold levels 20, 50 and 70%. We then compared the summary statistics between the three variant call format (VCF) SNP files. The VCF file with r=70% resulted in the least amount of missing data in the VCF and was kept for the subsequent analysis. We excluded sites with more than 50% missing data (option max-missing) from the resulting SNP VCF file by feeding it to VCFtools version 0.1.16 (Danecek et al., 2011). The amount of gDNA of the final set of samples ranged from 50 to 100ng. Samples that contained more DNA were more likely to pass the filtering steps (Fisher’s Exact Test for Count Data, p<0.05), but most samples passed these steps successfully regardless of the amount of DNA (50ng: 85%, 70ng: 89.4%, 100ng: 91.6%).

### Population structure and phylogeny analyses

We used several approaches to determine the population structure of *S. geminata*. Analyses were conducted in R version 3.5.0 (R Core Team, 2018). We conducted a Principal Component Analysis (PCA, snpgdsPCA function in the SNPRelate package, Zheng et al. 2012) based on our SNP data for 157 *S. geminata* samples to visualize how the samples from different locations clustered. To visualize clusters within the invasive populations, we also conducted a PCA excluding the native samples (i.e., Mexican) and another one without three additional samples (two from Ashmore Reef and one from India, Figure S1) because they clustered away from the remaining specimens and prevented us from looking at the structure of the invasive populations.

Phylogeny based on genome-wide SNPs was inferred on 156 individuals using IQ-TREE (Trifinopoulos et al., 2016). We uploaded the SNPs to the IQ-TREE web server and selected the automatic substitution model option and used the Ultrafast bootstrap analysis (Minh et al., 2013) with 1,000 bootstrap alignments to assess branch support. We used Dendroscope version 3.8.2 (Huson & Scornavacca, 2012) to visualise the output consensus tree file.

We used ADMIXTURE version 1.3.0 (Alexander et al., 2009) to estimate genetic ancestry on 155 individuals. For this analysis, we excluded the two most distant Mexican samples from the analysis based on the PCA results (Figure S2). ADMIXTURE uses a maximum likelihood approach to estimate the number of genetic clusters and the proportion of derived alleles in each sample from each of the K populations. We ran ADMIXTURE multiple times and varied the K value from 2 to 20. We used a cross-validation test to determine the optimal K value.

### Relatedness analysis

We used the populations program implemented in stacks (JM Catchen & Amores, 2011; Julian Catchen et al., 2013) to obtain the SNP data from 151 workers belonging to 22 colonies. We used r=1 to avoid biased estimation of relatedness from missing data (Kraemer & Gerlach, 2017). This step resulted in 84 to 922 loci per colony and 42 to 461 SNPs per colony. We estimated relatedness using Wang’s moment estimator of relatedness (Wang, 2002) implemented in Coancestry 1.0.1.9 (Wang, 2011). Wang’s relatedness estimator is more reliable than other indices for biallelic loci with allele frequencies of 0.5 (Wang, 2002). We estimated the effective mean number of queens (*N*_*qe*_) per colony as

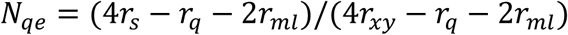

where *r*_*s*_ is the average relatedness of worker offspring from the same matriline, *r*_*q*_ the average relatedness of nestmate queens, *r*_*xy*_ the average relatedness of nestmate workers, and *r*_*ml*_ the average relatedness of the mates of nestmate queens (Ross, 1993; Seppä, 1994; Tay et al., 2011). We estimated *r*_*xy*_ from our data. We cannot determine *r*_*s*_, *r*_*q*_, and *r*_*ml*_ with our data and therefore relied on published data on polygyne *S. invicta* populations established in the USA to estimate their values. The invasive *S. invicta* is closely related to *S. geminata* and both species mate during nuptial flights (Hoffmann & O’Connor, 2004; Hung et al., 1977). The social structure of invasive *S. invicta* populations has been well-studied, whereas such information is lacking for invasive populations of *S. geminata* (but see Nipitwattanaphon et al., 2020 for an invasive population and Trible et al., 2018 for the native range).

We used *r*_*s*_ = 0.75 which is the average value for full sisters under male haploidy and outbreeding conditions (Crozier, 1970) as expected for *S. invicta*. We used *r*_*q*_ = 0 because queens in polygyne colonies of invasive *S. invicta* have a null average relatedness (Goodisman & Ross, 1997). Previous studies on the mating behaviour of *S. invicta* have found no evidence of local inbreeding (Ross, 1993; Ross et al., 1987; Ross & Fletcher, 1985). The mating flights of *S. invicta* and *S. geminata* involve alates from many colonies distributed over large areas which suggests panmixia (Lenancker et al., 2019). However, given the low genetic diversity of *S. geminata* (Gotzek et al., 2015; Lenancker et al., 2019), we expect some individuals from different colonies to be related. Therefore, we used two values of *r*_*ml*_, *r*_*ml*_ = 0 (mates of nestmate queens are unrelated) and *r*_*ml*_ = 0.5 (mates are brothers).

## RESULTS

### Data summary

From the 177 samples used to prepare the ddRAD libraries, we used a final sample set of 157 for the analysis after filtering the data (see SNP calling and filtering section in the methods) and obtained a total of 3,834 SNPs.

### Genetic bottleneck

Our results show that introduced *S. geminata* populations went through a strong genetic bottleneck across their genome. We obtained a total of 3,834 SNPs for all specimens vs only 2,358 SNPs for specimens collected in the introduced range. The PCA scores of specimens collected in the ant’s native range are interspersed among all the PCA scores (Figure S2) whereas the PCA scores of invasive samples form a compact group. Additionally, the number of SNPs in colony FC1 (461) from the native range exceeds that of any other colonies in the introduced range (average ± SD: 170.8±77.1, Table S2).

### Phylogenetic analysis

The phylogenetic analysis of the global *S. geminata* populations based on 156 specimens revealed a complex invasion history with multiple introduction events (Figure 2). Two strongly supported main clades suggest two main introduction waves: (i) Cambodia, the Philippines, Malaysia, Guam, and Australia (including mainland Australia, Christmas Island, Ashmore Reef, and Tiwi Islands) and (ii) Thailand, India, Hawai’i, and Micronesia.

**Figure 2.**
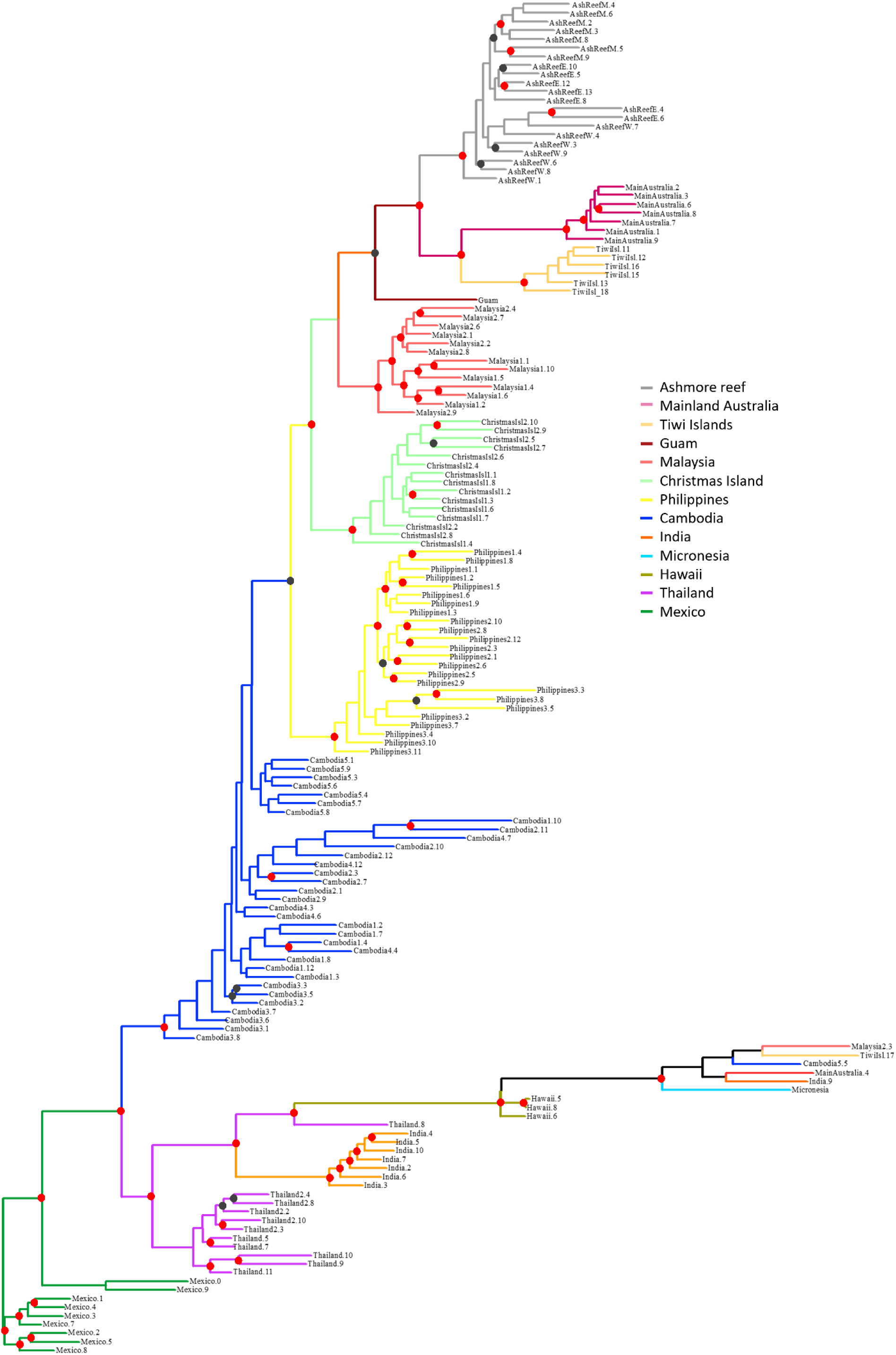
IQ-Tree with 1,000 bootstraps replications to estimate node support for *S. geminata* populations based on 156 specimens. Countries or islands are represented by different colours as indicated on the figure legend. Branch nodes with 85-100% bootstrap support are indicated by red dots and branch nodes with 74-84% support by grey dots. Colony code (Table 1) if there are more than one colony per location: AshReefM. M1A, AshReefE. E5A, AshReefW. W6A, MainAustralia.1 to MainAustralia.8 B1, MainAustralia.9 EF1.16PL, Malaysia1. DAG582.1, Malaysia2. SCCY14-07, ChristmasIsl1. Pink_House, ChristmasIsl2. Temple_Road, Philippines1. Phil1., Philippines2. Phil2., Philippines3. IRRI-700, Cambodia1. DAG621, Cambodia2. DAG623, Cambodia3. DAG618, Cambodia4. DAG622, Cambodia5. DAG637, Thailand. SH1, Thailand2 DAG561, Mexico.0 AVS2433, Mexico.9 Col1.1, and Mexico.1 to Mexico.8 FC1.

In the first main clade, the Cambodian population is ancestral to the Philippines, Malaysia, Guam, and Australia (74-84% bootstrap support). Tiwi Islands, mainland Australia, and Ashmore Reef form a well-supported clade (>85%), indicating a common introduction history for these populations. However, the phylogenetic analysis does not show which of these populations is ancestral to the others.

In the second main clade, Thailand is ancestral to India, Hawai’i, and Micronesia with strong bootstrap support (>85%). An individual from India and individuals belonging to populations in the first main clade (Malaysia, Tiwi Islands, Cambodia, Mainland Australia) form a well-supported clade potentially due to exchange of genetic material between the two main clades as shown in the ADMIXTURE analysis (Figure 3).

**Figure 3.**
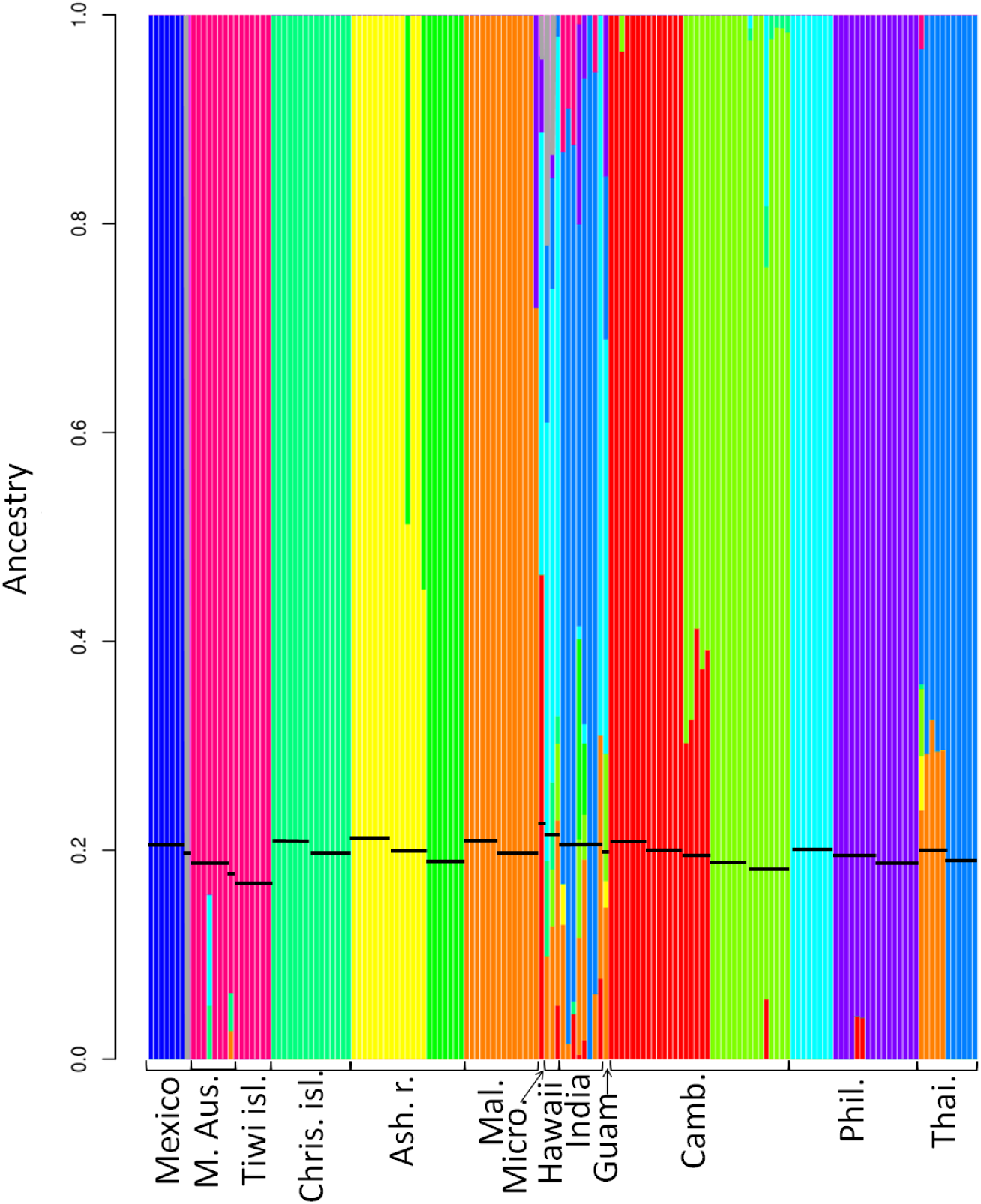
ADMIXTURE analysis conducted on SNPs from 155 specimens showing the optimal number of clusters: K=12. Individual black horizontal bars show specimen(s) belonging to the same colony. Abbreviations: M. Aus. mainland Australia, Tiwi isl. Tiwi Islands, Chris. isl Christmas Island, Ash. r. Ashmore Reef, Mal. Malaysia, Micro. Micronesia, Camb. Cambodia, Phil. The Philippines, Thai. Thailand.

### ADMIXTURE analysis

The ADMIXTURE analysis identified 12 genetic clusters (K=12) best describing *S. geminata* populations, confirming their complex population structure (Figure 3). Mexican specimens form two distinct clusters and are non-admixed.

The first main clade identified in the phylogenetic analysis (Cambodia, the Philippines, Malaysia, Guam, and Australia, Figure 2) forms nine clusters with or without evidence of gene flow from within the first clade, except for Guam for which there is evidence of gene flow from the second clade as well. Christmas Island forms an individual cluster with no evidence of gene flow from other clusters, whereas Malaysia also forms a distinct cluster but with evidence of gene flow from the Philippines. Mainland Australia and Tiwi Islands belong to the same cluster, confirming their common introduction history identified in the phylogenetic analysis (Figure 2). There is evidence of gene flow from Christmas Island, the Philippines, and Malaysia into mainland Australia. Cambodia forms two admixed clusters indicating movement of populations within the country. Cambodia also has evidence of gene flow from the Philippines and Ashmore Reef. The Philippines forms two clusters and one of them shows evidence of gene flow from mainland Australia/Tiwi Islands cluster.

Guam does not have its own cluster and is very admixed potentially explaining why it had poor support as ancestral to Tiwi Islands, mainland Australia, and Ashmore Reef in the phylogenetic analysis (Figure 2). Interestingly, Ashmore Reef forms two clusters despite the two most separated islands (West Island and East Island) only being approximately 13km apart, indicating that more than one introduction events took place. There is also evidence of gene flow from the West and East Island to the Middle Island as Middle Island belongs to a mix of the West and East Island clusters.

The second main clade identified in the phylogenetic analysis (Thailand, India, Hawai’i, and Micronesia, Figure 2) has mixed genomic signatures and none of them form a distinct cluster except for Thailand. Thailand shows evidence of gene flow from several populations in the first clade (e.g., Malaysia and Ashmore Reef). India, Hawai’i, and Micronesia have very mixed signatures from multiple populations in the first clade and Thailand. Interestingly, there is evidence of gene flow from one of the Mexican clusters to Hawai’i.

### Principal Component Analysis (PCA)

In the PCA analysis with 157 specimens including Mexico, the samples collected closest to Acapulco were the closest to the introduced cluster indicating that introduced populations potentially originated from this region (Figure S2 and Figure S3). The PCA with 144 specimens belonging to introduced populations shows that specimens from Ashmore Reef, Cambodia, and the Philippines each form distinct groups (Figure 4). Specimens from the other introduced populations are all mixed in one large group, albeit with individuals with the same geographic location tending to group closer together regardless of whether they belonged to the same clade identified in the phylogenetic analysis (Figure 2).

**Figure 4.**
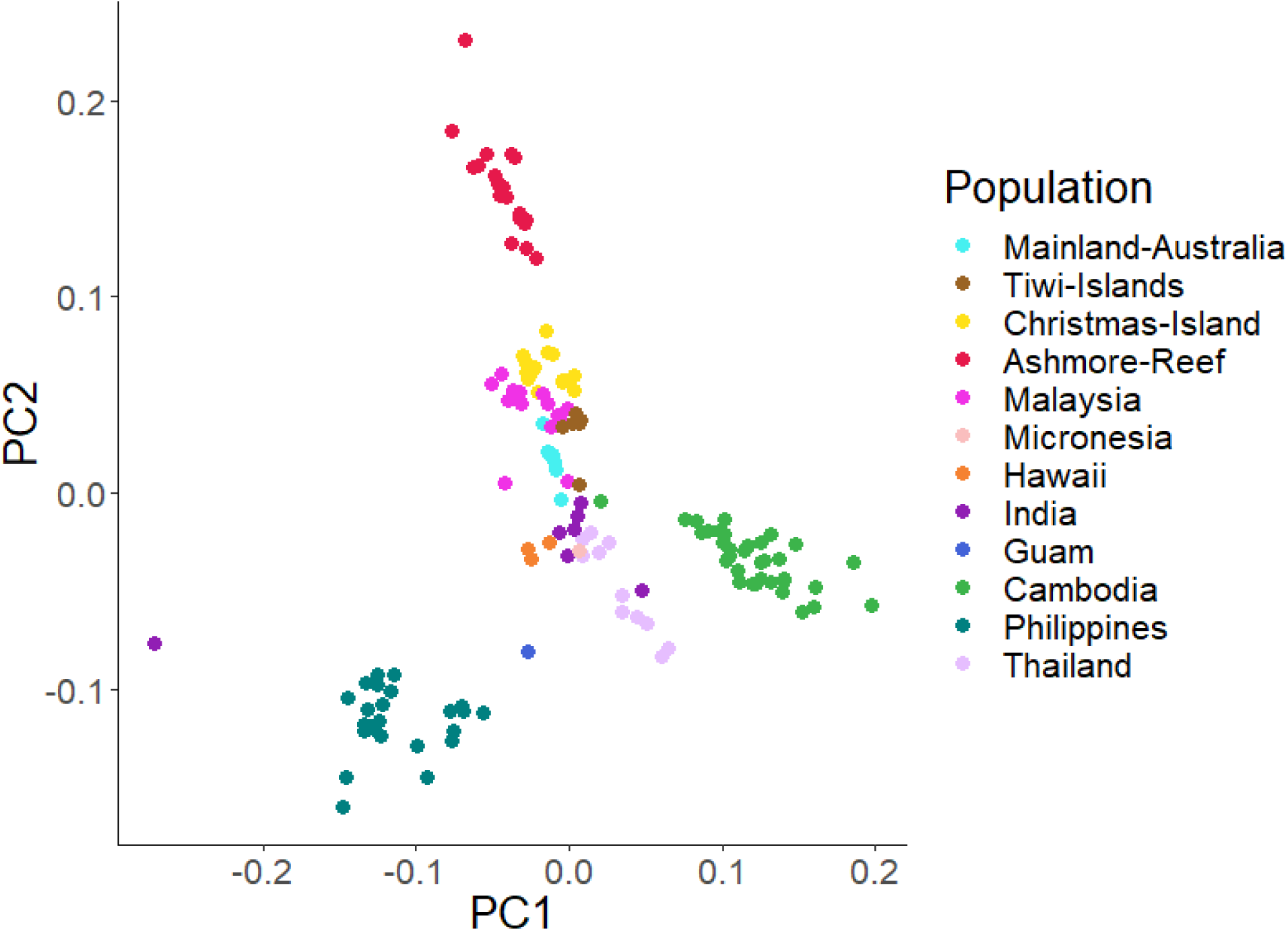
Principal component map of the SNPs in 144 individual specimens from the invasive range of *S. geminata*. The first axis represents 20% of the variation and the second axis 9%.

### Social structure

We determined the social structure for the 22 colonies, including one colony from the native range, for which we had more than one individual sampled. We found that all 22 colonies were polygyne (Table 2). The relatedness value *r*_*xy*_ (± SD) ranged from 0.26 ±0.13 to 0.68±0.11 (average ± SD: 0.12±0.05) which is lower than the expected relatedness value of 0.75 for monogyne colonies. We used an average (± SD) of 361.9±186.8 markers per colony for relatedness estimates. The effective number of queens (*N*_*qe*_) of polygyne colonies ranged from 1.1 to 40 queens per colony depending on the *r*_*ml*_ values (Table 2).

**Table 2.** Estimates of average nestmate workers relatedness (Wang’s r), effective number of queens (***N***_***qe***_) and social structure of colonies collected in the native and introduced ranges of *S. geminata*. We calculated ***N***_***qe***_ for two ***r***_***ml***_ (average relatedness of the mates of nestmate queens) values (see Table S2): *r*_*ml*_ = 0 (mates of nestmate queens are unrelated) and *r*_*ml*_ = 0.5 (mates are brothers) as we expect some individuals from different colonies to be related given the low genetic diversity of *S. geminata* in its introduced range (Gotzek et al., 2015; Lenancker et al., 2019). The lowest ***N***_***qe***_ value was calculated with *r*_*ml*_ = 0 and the highest with *r*_*ml*_ = 0.5 (Table S2).

| Sampling location | Colony code | Number of workers | Wang's $r$ ±SD | $N_{qe}$ | Social structure |
| --- | --- | --- | --- | --- | --- |
| mainland Australia | B1 | 7 | $0.68 \pm 0.12$ | 1.1-1.2 | polygyne |
| Tiwi Islands | Ti3 | 7 | $0.57 \pm 0.18$ | 1.3-1.6 | polygyne |
| Ashmore Reef | E5A | 7 | $0.54 \pm 0.15$ | 1.4-1.7 | polygyne |
| Ashmore Reef | M1A | 7 | $0.46 \pm 0.18$ | 1.6-2.4 | polygyne |
| Ashmore Reef | W6A | 7 | $0.53 \pm 0.08$ | 1.4-2.8 | polygyne |
| Christmas Island | Pink_House | 7 | $0.68 \pm 0.08$ | 1.1-1.2 | polygyne |
| Christmas Island | Temple_Road | 8 | $0.62 \pm 0.08$ | 1.2-1.3 | polygyne |
| Cambodia | DAG623 | 7 | $0.58 \pm 0.1$ | 1.3-1.5 | polygyne |
| Cambodia | DAG618 | 7 | $0.62 \pm 0.08$ | 1.2-1.4 | polygyne |
| Cambodia | DAG621 | 7 | $0.65 \pm 0.1$ | 1.2-1.3 | polygyne |
| Cambodia | DAG622 | 5 | $0.55 \pm 0.08$ | 1.4-1.6 | polygyne |
| Cambodia | DAG637 | 8 | $0.51 \pm 0.11$ | 1.5-1.9 | polygyne |
| Hawai'i | HDA | 3 | 0.26±0.13 | 2.9-40 | polygyne |
| India | HB-gem | 8 | 0.41±0.11 | 1.8-3.2 | polygyne |
| Malaysia | DAG582 | 6 | 0.61±0.08 | 1.2-1.4 | polygyne |
| Malaysia | SCCY14-07 | 8 | 0.68±0.11 | 1.1-1.2 | polygyne |
| Mexico | FC1 | 7 | 0.29±0.25 | 2.6-14.1 | polygyne |
| The Philippines | IRRI-700 | 8 | 0.52±0.11 | 1.4-1.8 | polygyne |
| The Philippines | Phil1 | 8 | 0.53±0.18 | 1.4-1.8 | polygyne |
| The Philippines | Phil2 | 8 | 0.61±0.04 | 1.2-1.4 | polygyne |
| Thailand | DAG561 | 5 | 0.54±0.07 | 1.4-1.7 | polygyne |
| Thailand | SH1 | 6 | 0.53±0.16 | 1.4-1.8 | polygyne |

## DISCUSSION

Our objectives were to (i) compare the genetic diversity of *S. geminata* in its introduced range to its most likely source of introduction, (ii) determine the invasion pathways among populations at an unprecedented level of detail for this species, and (iii) determine the social structure of *S. geminata* throughout its introduced range.

### Genetic bottleneck and inbreeding costs

Use of genome-wide SNP analysis enabled us to determine that *S. geminata* populations went through a strong genetic bottleneck across their genome following introduction events, which may have repercussions for colony fitness. A total of 3,834 SNPs was found for all specimens vs only 2,358 SNPs for specimens collected in the introduced range. We consider this a minimum estimate of the loss of genetic diversity because our study only included specimens collected from the most likely source of introduced populations (i.e., southwestern Mexico, Gotzek et al., 2015). In previous research, introduced populations of *S. geminata* were found to have 93% fewer *mtCOI* haplotypes and 63% fewer microsatellite allele on average than native populations (Gotzek et al., 2015). In many invasive species, introduced populations go through a genetic bottleneck as they establish from a small number of founder individuals (Allendorf & Lundquist, 2003). As a result, their genetic diversity is often a fraction of native populations’ diversity (Allendorf & Lundquist, 2003; Dlugosch & Parker, 2008). Loss of genetic diversity between native and introduced ant populations is also common and can generate severe consequences on the colony growth of some species or may contribute to their success (Ascunce et al., 2011; Lenancker et al., 2019; Ross et al., 1993; Tsutsui & Suarez, 2003; Vogel et al., 2010).

Loss of genetic diversity can generate severe consequences on the colony growth of invasive ants as shown for *S. invicta* and *S. geminata* (Lenancker et al., 2019; Ross & Fletcher, 1986). Low genetic diversity can lead to the loss of CSD (complementary sex determination) alleles which disrupts the functioning of the sex determination system (Ross et al., 1993). Generally in the Hymenoptera, when a queen and the male she mated with share the same CSD genotype, half of the queen’s offspring resulting from this mating will develop into diploid males instead of workers (Crozier, 1971, 1977; Heimpel & de Boer, 2008). Adult diploid males do not contribute to the reproductive output of the colony or its workload (Cook & Crozier, 1995; Ross & Fletcher, 1986). In *S. geminata* and *S. invicta* diploid male production can reduce colony growth and survival (Cook & Crozier, 1995; Lenancker et al., 2019; Ross & Fletcher, 1986). Diploid male production is common in Australian *S. geminata* colonies but strategies during colony founding, such as pleometrosis (queens founding their nest together) and execution of diploid male larvae, can minimize its effects during colony founding (Lenancker et al., 2019). Another invasive ant, *Brachyponera chinensis*, was found to be pre-adapted to potential inbreeding depression (Eyer et al., 2018). Both native and introduced populations have low genetic diversity because *B. chinensis* queens preferably mate with their brothers. Sibmating in the native range may reduce the potential cost of inbreeding for *B. chinensis* by purging deleterious alleles contributing to their success as invaders (Eyer et al., 2018).

For *L. humile*, loss of genetic diversity may have contributed to its invasive success (Tsutsui & Suarez, 2003). In their native range, *L. humile* workers can differentiate nestmates from outsiders because different colonies have distinct cuticular hydrocarbon profiles (Brandt et al., 2009). In its invasive range, however, *L. humile* is unicolonial, presumably because the cuticular hydrocarbons of introduced populations became homogeneous as a consequence of genetic bottlenecks (Brandt et al., 2009; Tsutsui et al., 2000). In the absence of costs related to intraspecific territoriality, colonies in the introduced range are potentially able to direct more resources toward foraging, colony growth, and interspecific competition than colonies in the native range (Tsutsui & Suarez, 2003).

### Invasion pathways

The bridgehead effect is a major driver increasing global rates of ant invasions (Bertelsmeier et al., 2018), and the use of genome-wide SNP data analyses in our study reveals the complex population movements occurring between introduced *S. geminata* populations at an unprecedented level of detail. A previous study using microsatellite and mitochondrial markers to identify global invasion routes suggested that *S. geminata* was accidentally introduced to the Old World by Spanish galleons departing from Acapulco to Manila (the Philippines) and Canton (China) from the 16^th^ to the 18^th^ century (Bjork, 1998; Flynn & Giráldez, 1995; Fradera, 2004; Gotzek et al., 2015). However, before now detailed movements among introduced populations of *S. geminata* could not be determined due to the low variability of markers across its introduced range. We found that the closer to Acapulco the ants were collected, the closer they were to introduced populations on the PCA (Figure S2 and S3). Populations established in the Philippines were related to some, but not all invasive populations, revealing a more complex history than previously described (Gotzek et al., 2015). We found that invasive populations originate from two main locations: Cambodia and Thailand (Figure 2). The Cambodian population invaded the Philippines, Malaysia, Guam, and Australia. The Thailand population invaded India, Hawaii, and Micronesia. We also found that some individuals from Malaysia, Australia, and Cambodia were closely related to the populations originating from Thailand indicating movement among the populations originating from Cambodia and Thailand. Overall, our results indicate that secondary invasions from multiple locations have occurred confirming bridgehead effect is a driving factor in the establishment of global ant invaders (Bertelsmeier et al., 2018).

Additionally, we identified a high number of gene flow events among invasive populations indicating that colonies of *S. geminata* have regularly been transported among these locations. For example, we found evidence of gene flow going from the Australian mainland to India and from Malaysia to Thailand. These population movements are potentially still taking place. Connection among invasive populations and multiple introductions can be problematic because they increase the genetic diversity in invasive populations and reduce the deleterious effects of inbreeding in the long term (Dlugosch & Parker, 2008). Continuous gene flow among invasive *S. geminata* populations may lead to an increase in the diversity of CSD alleles and decrease the occurrence of diploid male production in some populations over a long period of time, potentially increasing their colony growth and spread.

Our results have also revealed the common introduction history of Australian populations (i.e., mainland Australia, Tiwi Islands, Christmas Island, and Ashmore Reef) which all show a Cambodian origin. Christmas Island may be an independent invasion event to other Australian populations due to clustering away from the other Australian populations. *Solenopsis geminata* was first recorded on Christmas Island in 1915, after mainland Australia’s first record in Darwin (Northern Territory) prior to 1863 (Wetterer, 2011) indicating that Darwin may have been colonised by *S. geminata* prior to Christmas Island. However, this first record for mainland Australia may not be accurate as we did not find location information in the literature cited (Roger, 1863) by Wetterer (2011), and Darwin was not settled by Europeans until 1869 (Taylor & Lea, 1988). We found evidence of gene flow from Ashmore Reef to several other populations (e.g., Thailand and India, Figure 3). Ashmore Reef is currently uninhabited and consists of three main islands which are 320km off the Australian northwest coast and 144km south of Rote Island in Indonesia.

Despite their remoteness, guano was mined from the islands during the 19th century, and fishermen from Indonesia have frequently visited them for 300 years, potentially displacing ants to and from Ashmore Reef by accident (Clarke et al., 2011; Hodgson & Clarke, 2014). We also cannot exclude that Ashmore Reef acts as a proxy population for other potential source populations and that the gene flow events from Ashmore Reef may in fact be attributed to other populations such as Indonesia, from which we were unable to obtain specimens.

Our results highlight the importance of using a combination of genomic tools and occurrence data to determine the introduction pathways of invasive species. Retracing the invasion pathways and native range using occurrence records is difficult for historically introduced ant invaders such as *S. geminata*, and using genomic tools may be the only option to resolve uncertainties in their invasion history. For example, Caribbean populations of *W. auropunctata* were proven to be introduced and not native to the area by a genetic study (Foucaud et al., 2010). Additionally, despite being listed among the world worst’ invaders, the native range of *Anoplolepis gracilipes* (yellow crazy ants) remains unknown although the relatively high mitochondrial DNA haplotype diversity of populations in southeast Asia suggests that it originated from this region (Drescher et al., 2007; Wetterer, 2005). Retracing invasion pathways of recent invaders using detection data may also prove challenging. For example, initial detection of *Spodoptera frugiperda* (fall armyworm) in West Africa in 2016 followed by detection across the Old World supported an eastward expansion of this species (Tay et al., 2022). However, genomic evidence has shown that populations of *S. frugiperda* found in Africa originate from Asia (Tay et al., 2022). Despite the unprecedented level of detail provided by RADseq data analysis, we did not distinguish historical from recent invasion events, which could potentially be useful information for biosecurity management plans. Using a combination of genomic, historical occurrence, and interception data may be the best option to clarify population movements of invaders and may provide useful information for targeted biosecurity management programs.

### Social structure of *S. geminata* across its invasive range

We found that all colonies in the invasive range were polygyne which may have consequences for the invasion success and dispersal of *S. geminata*. Polygyny can increase colony growth rate and the establishment potential of transported colony fragments because they are more likely to contain a queen (Tsutsui & Suarez, 2003; Vargo & Fletcher, 1989). Polygyne populations of invasive ants can also reach higher densities than monogyne ones, which contributes to their ability to dominate ecosystems (Holway et al., 2002; Macom & Porter, 1996). In Mexico, polygyne *S. geminata* nest density was estimated to be more than 40 times higher than nest density of an adjacent monogyne population (Mackay et al., 1990). Whether an invasive ant population is monogyne or polygyne may also affect management strategies. For example, most *S. invicta* colonies in southeast Queensland (Australia) are monogyne and spread by flight, whereas polygyne colonies spread through budding or movements of materials (Wylie et al., 2016). As a result, surveillance buffers for monogyne colonies are larger than for polygyne colonies to account for new colonies founded by flying queens (Wylie et al., 2016). Interestingly, it was recently revealed that most queens in polygyne colonies from a *S. geminata* population in Florida were produced asexually and workers sexually, whereas both castes were produced sexually in monogyne colonies (Lacy et al., 2019). Additional investigations into the reproductive systems of *S. geminata* will reveal whether this mixed reproductive system associated with colony social structure is present in other populations and its role in the invasion success of *S. geminata*.

## Conclusions

The use of ddRADseq enabled us to use more than 3,500 SNPs to compare the genetic diversity of *S. geminata* in its introduced range to its most likely source of introduction, determine the invasion pathways among introduced populations in unprecedented detail, and determine the social structure of *S. geminata* throughout its introduced range. This technique is ideal for small taxa as samples with as little as 50ng of DNA passed the filtering steps with a high success rate (85%). Our findings highlight the importance of bridgehead effect as a major driver of the spread of *S. geminata* with multiple introductions occurring among populations in the introduced range.

## Acknowledgements

We are grateful to our research partners and institutions who collected and shared samples for this study: J. Amith, H. Bharti, R. M. Clouse, L. Cruz-López, F. Cupul, M. Fukada, S. Hasin, A. Herrod, B. Howes, D. Maple, P. Pantaleón, A. V. Suarez, S. Yang, and the International Rice Research Institute. This work was supported by a Holsworth Wildlife Research Endowment from the Ecological Society of Australia, awarded to PL.

## Supplementary information

**Table S1.**
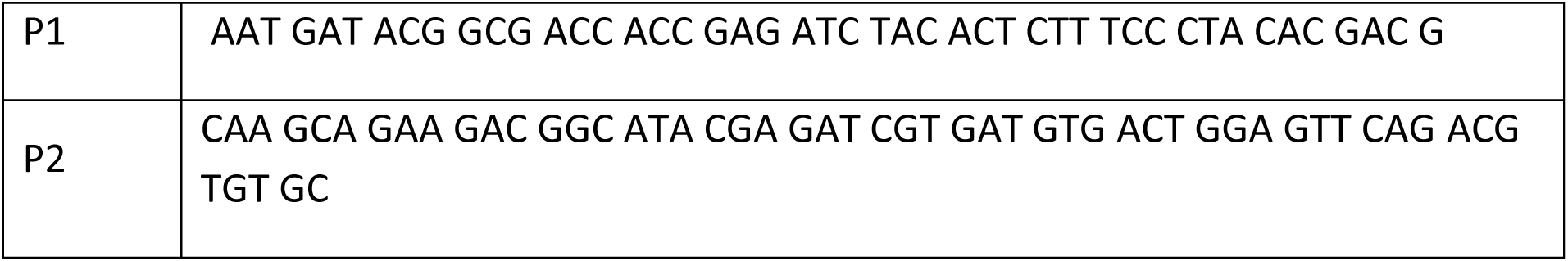
Oligo sequence of primer pairs for PCR reaction

**Table S2.** Number of loci for each colony in which multiple nestmates were collected, estimates of average relatedness of nestmate workers (Wang’s r), effective number of queens (***N***_***qe***_) for two ***r***_***ml***_ (average relatedness of the mates of nestmate queens) values.

| Origin | Colony<br>code | SNPs/loci | Wang's $r \pm SD$ | $N_{qe}$ | |
| --- | --- | --- | --- | --- | --- |
| | | | | $r_{ml}=0$ | $r_{ml}=0.5$ |
| Mexico | FC1 | 461/922 | 0.29±0.25 | 2.6 | 14.1 |
| Mainland<br>Australia | B1 | 54/108 | 0.68±0.12 | 1.1 | 1.2 |
| Tiwi Islands | Ti3 | 79/158 | 0.57±0.18 | 1.3 | 1.6 |
| Christmas<br>Island | Pink_House | 202/404 | 0.68±0.08 | 1.1 | 1.2 |
| Christmas<br>Island | Temple_Road | 169/338 | 0.62±0.08 | 1.2 | 1.3 |
| Ashmore<br>Reef | E5A | 164/396 | 0.54±0.15 | 1.4 | 1.7 |
| Ashmore<br>Reef | M1A | 298/396 | 0.46±0.18 | 1.6 | 2.4 |
| Ashmore<br>Reef | W6A | 241/482 | 0.53±0.08 | 1.4 | 2.8 |
| Malaysia | DAG582 | 178/356 | 0.61±0.08 | 1.2 | 1.4 |
| Malaysia | SCCY14-07 | 65/130 | 0.68±0.11 | 1.1 | 1.2 |
| Hawai'i | HDA | 198/396 | 0.26±0.13 | 2.9 | 40 |
| India | HB-gem | 60/120 | 0.41±0.11 | 1.8 | 3.2 |
| Cambodia | DAG623 | 126/251 | 0.58±0.1 | 1.3 | 1.5 |
| Cambodia | DAG618 | 263/526 | 0.62±0.08 | 1.2 | 1.4 |
| Cambodia | DAG621 | 151/302 | 0.65±0.1 | 1.2 | 1.3 |
| Cambodia | DAG622 | 207/414 | 0.55±0.08 | 1.4 | 1.6 |
| Cambodia | DAG637 | 42/84 | $0.51 \pm 0.11$ | 1.5 | 1.9 |
| Philippines | IRRI-700 | 187/374 | $0.52 \pm 0.11$ | 1.4 | 1.8 |
| Philippines | Phil1 | 194/388 | $0.53 \pm 0.18$ | 1.4 | 1.8 |
| Philippines | Phil2 | 213/426 | $0.61 \pm 0.04$ | 1.2 | 1.4 |
| Thailand | DAG561 | 329/657 | $0.54 \pm 0.07$ | 1.4 | 1.7 |
| Thailand | SH1 | 167/334 | $0.53 \pm 0.16$ | 1.4 | 1.8 |

**Figure S1.**
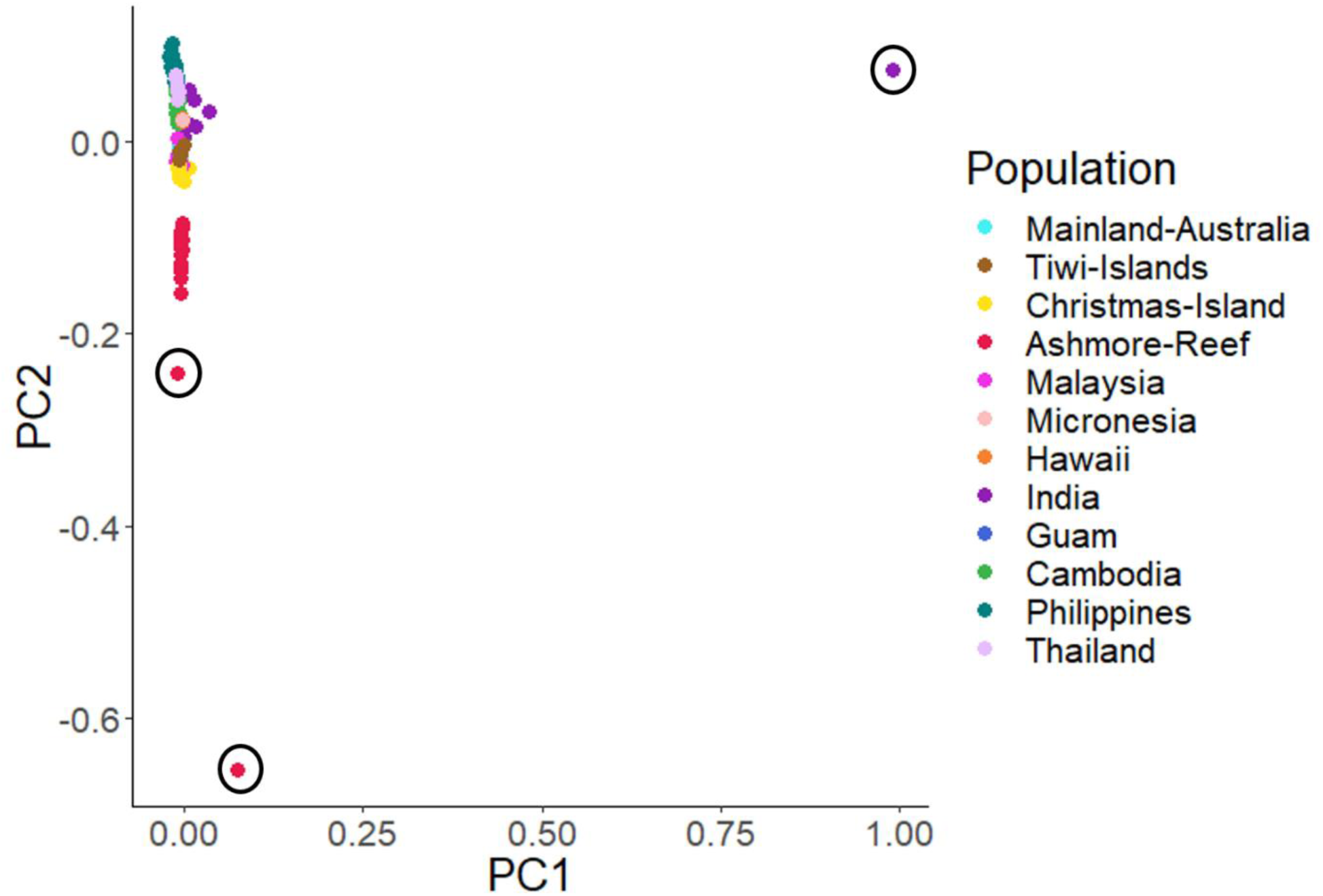
Principal component map of the SNPs in 147 individual specimens from the invasive range of *S. geminata*. We removed the circled specimens from the subsequent principal component map (Figure 4) to observe how the other invasive specimens clustered together.

**Figure S2.**
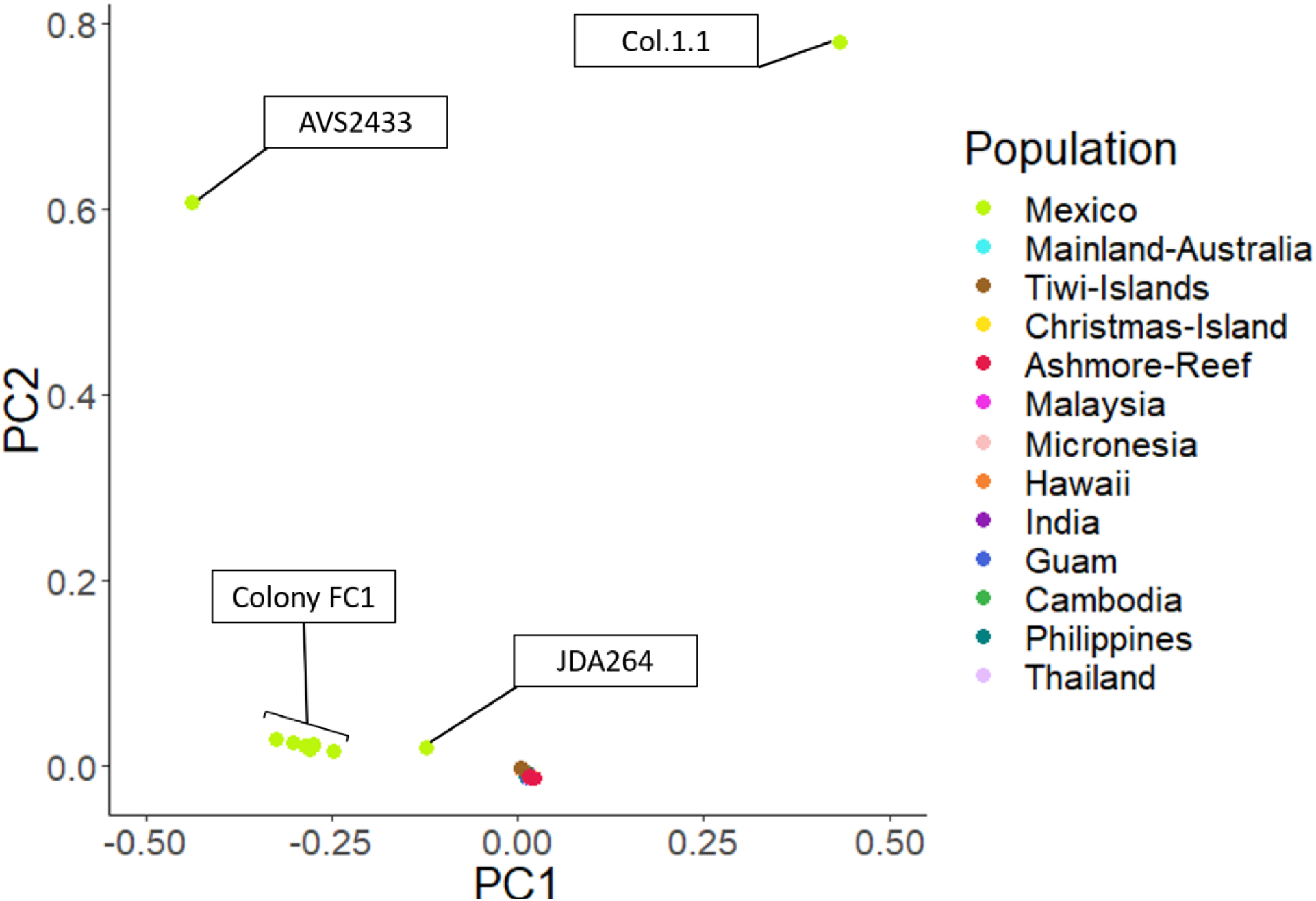
Principal component map of the SNPs in 157 individual specimens. The specimens originating from Mexico are labelled with their individual code or their colony code (Table 1). See Figure S3 for a map showing the origin of the Mexican specimens.

**Figure S3.**
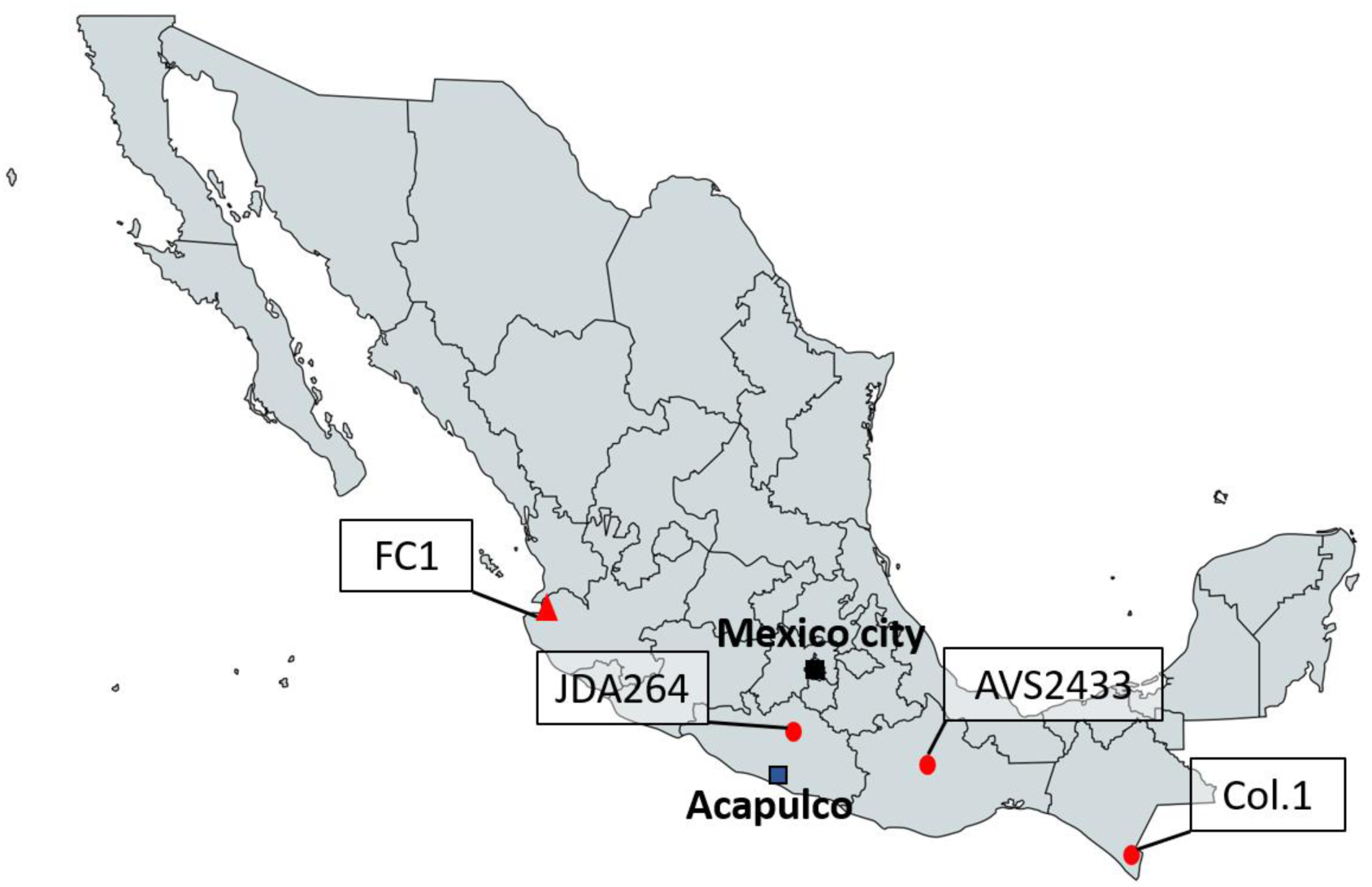
Map of the Mexican’s sample origin with the sample codes. Circle: single specimen. Triangle: multiple specimens from a colony.

